# Experimentally assessing the role of foraging encounters on social partner choice in a kin-based bird society

**DOI:** 10.1101/2025.11.03.686260

**Authors:** Babette Fourie, Claire Doutrelant, Yoann Depalle, Andre C. Ferreira, Rita Covas

**Affiliations:** CIBIO, Centro de Investigação em Biodiversidade e Recursos Genéticos, InBIO Laboratório Associado, Campus de Vairão, Universidade do Porto, 4485-661 Vairão, Portugal; BIOPOLIS Program in Genomics, Biodiversity and Land Planning, CIBIO, Campus de Vairão, 4485-661 Vairão, Portugal; CEFE, Univ Montpellier, CNRS, EPHE, IRD, Montpellier, France; FitzPatrick Institute of African Ornithology, DST-NRF Centre of Excellence, University of Cape Town, Rondebosch 7701, South Africa; Department of Evolutionary Biology and Environmental Studies, University of Zurich, Zurich 8057, Switzerland

## Abstract

Kin-based social environments are often characterized by prolonged parent-offspring associations and limited dispersal in at least one sex, leading to groups composed of kin as social partners. These high rates of social interactions among relatives in cooperative societies make it difficult to distinguish whether social partner choice is driven by kinship or by repeated encounters with individuals situated at proximity. Here, we experimentally increased the number of foraging encounters among non-kin at a custom feeder setup in the sociable weaver (Philetairus *socius*), a colonial, cooperatively breeding bird with a kin-based social structure. Individuals encouraged to forage more frequently with non-relatives subsequently developed stronger foraging associations with non-kin at separate, unrestricted feeders. These findings show that repeated opportunities for interaction can promote non-kin social associations even within a kin-based society, contributing to the understanding of proximate mechanisms that underlie social structure.

## Introduction

Individuals in kin-structured populations are typically born into and remain near kin during development, and hence disentangling the effects of active kin preference in social associations from spatial proximity becomes challenging (Riehl & Strong., 2018; Greenwood, 1980; Clutton-Brock & Lukas, 2011). Inclusive fitness theory (Hamilton, 1964) predicts that individuals can gain indirect fitness benefits through associating and cooperating with kin (Maynard Smith, 1964), in addition to any possible direct fitness benefits (Clutton-Brock, 2009), providing a powerful explanation for the evolution of social behaviour. Such indirect fitness benefits can arise if individuals actively select to associate and cooperate with kin (i.e. discriminate altruism; Taborsky et al., 2021) or associate and cooperate with kin as a result of spatial overlap with relatives (i.e. population viscosity leading to indiscriminate altruism: Mitteldorf & Wilson., 2000; Hamilton, 1964).

This distinction is relevant because observations of kin-based associations are often interpreted to be evidence for preference to help kin, and thus of indirect fitness benefits (e.g. Dickinson & Hatchwell., 2004). In many species, individuals also choose to associate with non-kin. For example, vampire bats (*Desmodus rotundus)* engage in reciprocal food sharing with hungry social partners, including both non-relatives and relatives (Carter & Wilkinson., 2013) and in Assamese macaques (*Macaca assamensis)*, where individuals form enduring stable relationships, or *“*social bonds”, that affect fitness, males had similar bonding propensity with kin and non-kin males (De Moor et al., 2020). Similarly, in approximately 30% of cooperatively breeding bird species, there is a mix of related and unrelated helpers (Riehl, 2013). These studies suggest that there might be direct fitness benefits for cooperating with social partners, not dependent on kinship. However, in many species, because of limited dispersal and/or the social environment animals are born into, it might be easier to form partnerships with kin. This can lead to an intertwine between kinship and social partnerships and thus to potentially an overemphasis of the role of indirect fitness benefits.

Experimentally investigating the proximate mechanisms underlying social bond formation, particularly in kin-based societies, can help us to understand social partner choice, and whether associating with and helping kin or non-kin ultimately constitutes evidence for indirect fitness benefits, kin discrimination errors (Morinay et al., 2025) or direct benefits (Carter & Wilkinson., 2013). Such experiments can also reveal how population (Mittledorf & Wilson., 2000) and social (Grabowska-Zhang et al., 2016) viscosity limit the options for social partner choice. In kin-based societies, some previous studies relied on cross-fostering experiments to investigate the proximate causes of social associations (Sharp et al., 2005). An alternative more natural approach is to study foraging interactions of individuals during adulthood, where repeated encounters can promote and maintain social affiliations that might eventually lead to bond formation and cooperative interactions.

Sociable weavers (*Philetairus socius*) are an excellent study system to test whether social partner choice is shaped by repeated foraging opportunities to associate. They are colonial cooperative breeders that form kin-based associations for breeding (Covas et al., 2006; Ferreira et al., 2024), roosting (Paquet et al., 2016), and foraging (Ferreira et al., 2020). Most helpers at the nest are kin (Covas et al., 2005; Ferreira et al. in prep) and form social bonds with breeders, which can be detected during foraging (Ferreira et al., 2020). Foraging offers a key context for social bond formation, providing benefits such as improved information access and reduced predation risk (Lima., 1995; Valone & Templeton., 2002; Berdahl et al., 2013;). We designed a new experimental method to test whether artificially increasing foraging encounters between non-kin leads to the formation and strengthening of non-kin social associations (Figure 1). The study was conducted on three wild colonies of sociable weavers at a long-term study site in Benfontein Nature Reserve, South Africa (Covas et al., 2006).

**Figure 1.**
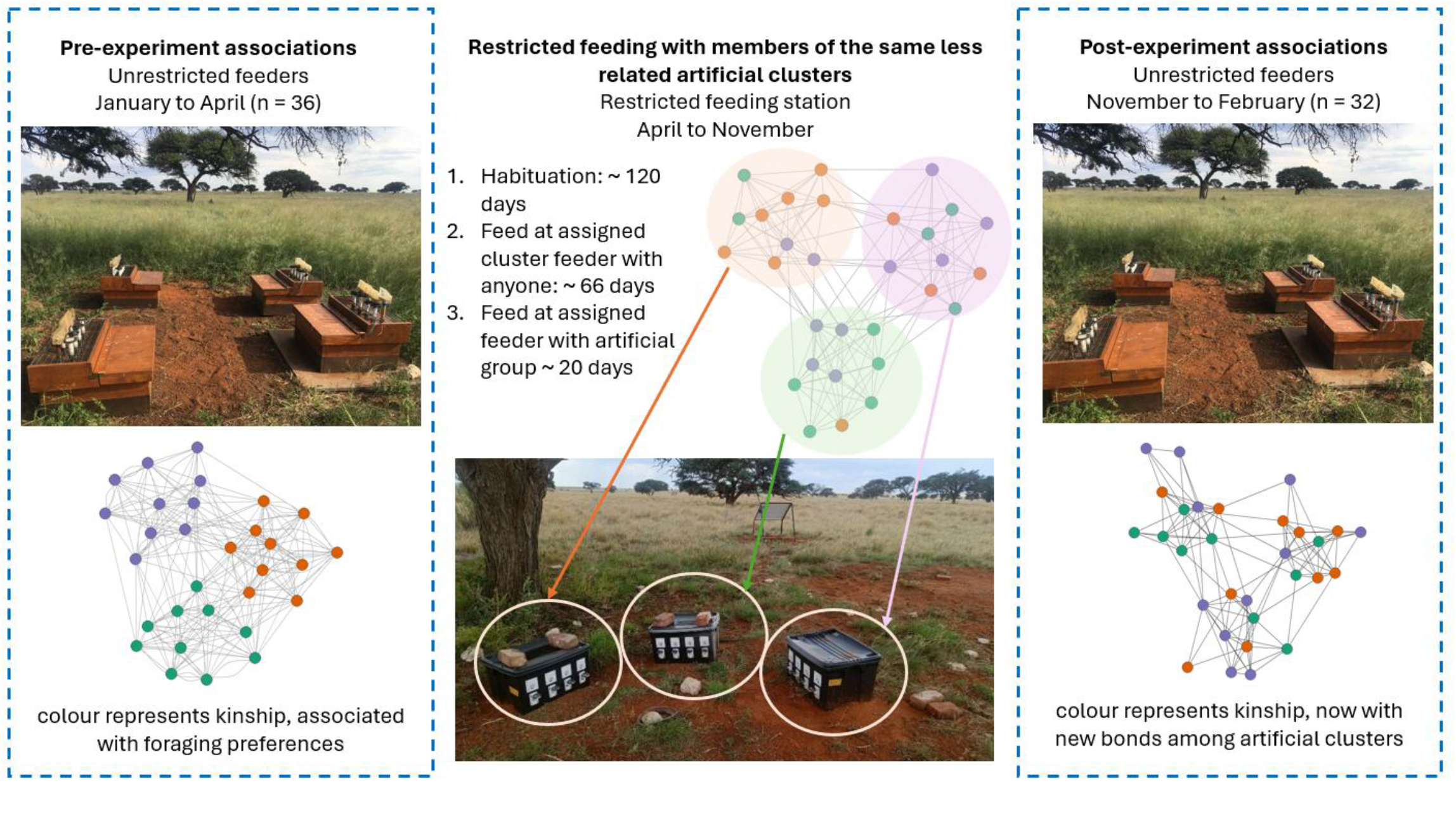
Three stages of the experiment: (A) Pre-manipulation social foraging associations were recorded at unrestricted feeders, showing natural kin-based preferences. (B) Individuals were grouped into artificial clusters with low genetic relatedness, and consequently lower co-foraging preferences, and given access to restrictive feeders controlled by RFID technology, allowing only artificial cluster members to feed together. (C) Post-manipulation associations from unrestricted feeders showed an increase in associations within artificial clusters. Each of the three sociable weaver colonies had access to one unrestricted and one restricted feeding station at separate locations.

Using RFID technology (Radio Frequency Identification), we designed selective feeders that restricted food access among kin (i.e., preferred foraging partners; Ferreira et al., 2020) and encouraged less-related individuals (typically less preferred foraging partners) to feed together (Figure 1). We created experimental groups, by combining genetic relatedness data with clustering algorithms (Louvain & hierarchical clustering, see Supplementary Information) to divide individuals into artificial clusters that minimized within-group relatedness. Each cluster of less-related individuals was assigned to a single restrictive feeder within a station of three or four feeders (Figure 1). After a period of habituation (May 2024 – October 2024), we conducted a restrictive manipulation phase (November) during which individuals from each cluster could feed either alone or with other less-related members of their assigned group (SI appendix). We assessed the effect of this manipulation by constructing time-overlap social networks (Ferreira et al., 2020) from foraging associations recorded at spatially separate, unrestricted feeders both before (January 2024 -April 2024) and after the manipulation (November 2024-February 2025; Figure 1).

## Results and Discussion

For each sociable weaver colony, individuals were assigned to artificial clusters that had significantly lower relatedness than natural foraging groups (Welch’s t-test: t = –3.44, p < 0.05, 95% CI [–0.20, –0.04]; mean related edge proportion: artificial = 0.043, natural = 0.165). Dyads within these artificial clusters were allowed to feed together at the manipulation feeders, and dyads from different clusters could not feed together.

During the manipulation at the restrictive feeders, assortativity by artificial cluster increased in all three colonies, shifting from negative values before the manipulation to positive values during the manipulation (Colony 8, r = -0.019 to 0.257; Colony 11, r = -0.127 to 0.105; Colony 71, r = -0.119 to 0.135). This confirmed that the manipulation successfully increased within-cluster foraging associations.

Before the experiment, within-cluster dyads had lower association strengths than the between cluster dyads (β = −0.75, 95% CrI [−1.22, −0.32]); Figure 2). This indicates that individuals assigned to the same artificial clusters had weaker foraging associations before the manipulation. Between cluster dyads declined from pre-to post-manipulation (β = −0.76, 95% CrI [−0.88, −0.63]), whereas within-cluster dyads that had encounters at the manipulation feeders showed an increase in association strengths at the unrestricted (post-experiment) feeders relative to between cluster dyads (interaction β = 1.01, 95% CrI [0.46, 1.57]; Figure 2). Model-predicted association strengths showed that between cluster dyads decreased from 0.137 to 0.072, whereas treatment dyads increased from 0.074 to 0.092.

**Figure 2.**
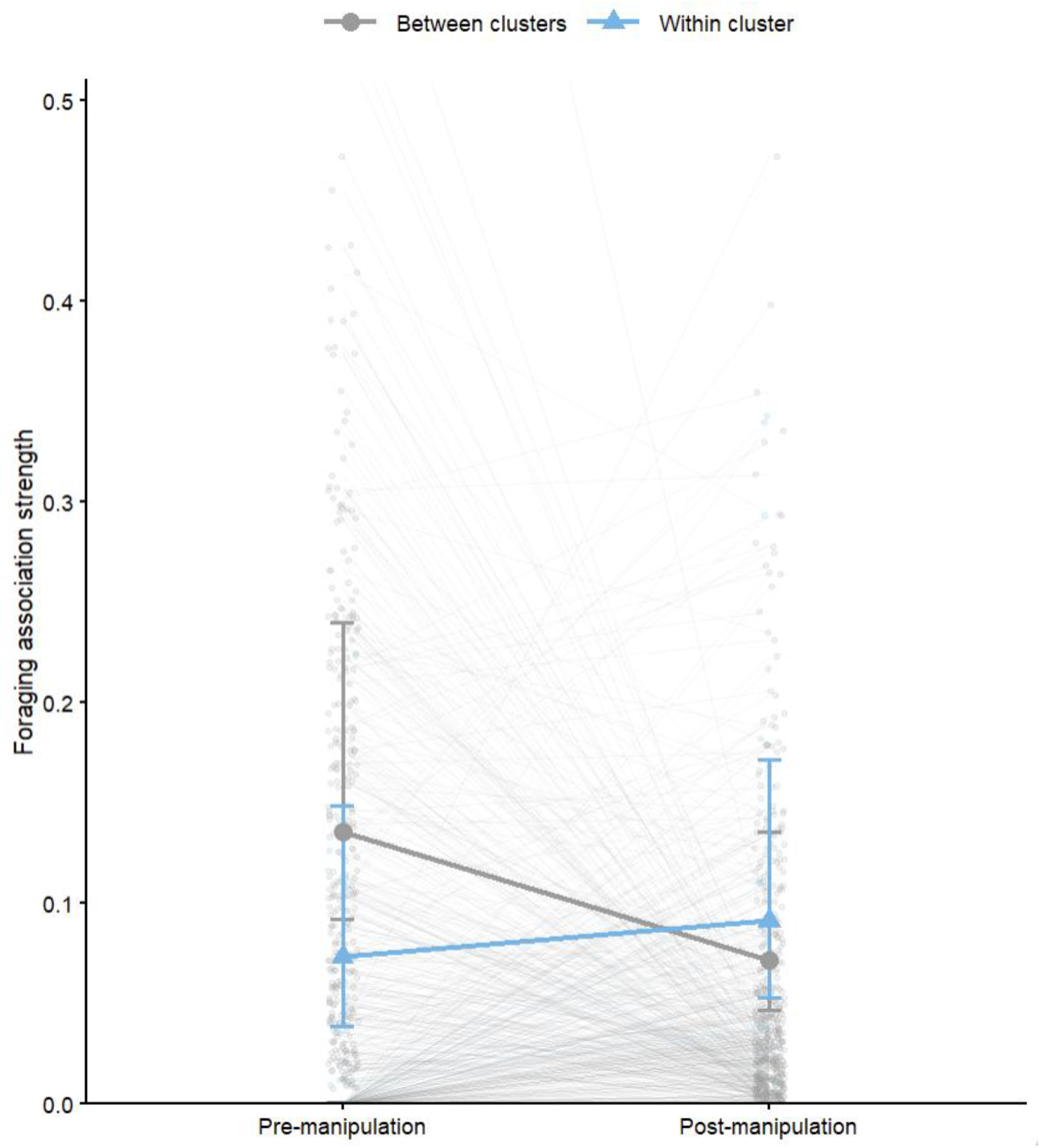
Changes in foraging association strength before and after the manipulation. Model predicted mean association strengths and 95% Bayesian credible intervals are shown for between-cluster dyads (grey; pre = 0.137, post = 0.072) and within-cluster dyads (blue; pre = 0.074, post = 0.092). Thin background lines connect observed association strengths for individual dyads before and after the manipulation. Analyses were restricted to within-cluster dyads that frequently used the restrictive feeders, and between-cluster dyads for which both individuals were observed at the restrictive feeders.

These results demonstrate that foraging associations increased among less-related individuals that fed together during the manipulation, supporting our prediction that repeated interactions can foster (at least short-term) non-kin associations, even in a kin-based social system. Hence, our results demonstrate that social partner choice remains flexible despite strong kin structuring.

Previous experimental manipulation of social associations through conditional access to food has successfully altered short-term foraging associations in other species and contexts (Firth et al., 2015; King et al., 2023; Heinen et al., 2022), showing that individuals have the capacity to adjust social partner choice when needed. However, to our knowledge, our study provides the first experimental demonstration that repeated foraging encounters alone can strengthen non-kin social associations in a kin structured cooperative society.

Evidence of this flexibility in social partner choice is particularly important in societies with kin-based associations patterns, where opportunities for social interactions are often confounded with kinship (i.e., active kin discrimination and preference). Consequently, this overlap can have important implications for interpreting what constitutes evidence for kin selection: apparent preferences for helping kin over non-kin (or closer over more distant relatives) may reflect spatial and social network constraints in social association development as much as shared genes. This may complicate interpretations of the relative roles of direct and indirect fitness benefits (Garcia-Ruiz et al., 2022), highlighting the need for further empirical and experimental investigation into how the development and strength of social associations shape cooperative decisions.

## Acknowledgments

We thank Franck Theron and all field workers who helped with electronics and RFID data collection, particularly Antoine Grissot, Leo Jhaveri, Irene Baquero, Cristina Jimenez, and Hippolyte Dupas. Thanks to Damien Farine for his comments on the manuscript. De Beers Consolidated Mines provided access to Benfontein Reserve. This study was funded by an ERC Consolidator grant 866489 (EU) to RC; OSU OREME and SEE-Life (CNRS, France) to CD; FCT (CEECIND/03451/2018) to RC.

## Supplementary Information

### Methods

Access to data and code is here https://doi.org/10.6084/m9.figshare.29775938.v1

#### (A) Study system

This experiment was conducted as part of a long-term research program on a population of social weaver, *Philetairus socius*, colonies at Benfontein Nature Reserve, near Kimberley, South Africa (Covas et al., 2006). Sociable weavers are cooperatively breeding passerines that live year-round in large communal nests composed of multiple, independent chambers. These nests are used as breeding sites during the reproductive season and as roosting sites year-round. Colonies in this population typically range from 10 to 120 individuals and can contain up to 78 different nest chambers.

For this study, we manipulated three sociable weaver colonies, respectively named “8, 11, and 71”. These colonies were previously fitted with metal rings and unique colour combinations, including PIT-tags (Passive Integrated Transponder) integrated into leg rings. Individuals are captured annually to be ringed, and blood sampled for sex determination and genotyping (Paquet et al. 2015). During the captures before the experiment (August 2023), the three colonies had the following number of recorded individuals: Colony 8 = 84 individuals, Colony 11 = 53 individuals, Colony 71 = 38 individuals.

Sociable weavers exhibit variable lengths of breeding periods (D’Amelio et al., 2024). This experiment was conducted throughout 2024, with the pre-experiment phase (January to April) corresponding to the end of the 2023/2024 breeding period, the habituation phase (May to November) occurring during the non-breeding period, and the post-experiment period (December 2024) overlapped with the start of the 2024/2025 breeding period.

#### (B) Experiment overview and artificial clusters

We aimed to encourage foraging associations between less related individuals and, consequently, restrict their preferred kin-based foraging associations. We did this by using restrictive feeders that facilitated feeding interactions between artificially created groups of less-related individuals. The restrictive feeders allowed individuals to gain food access either alone or alongside to their artificial group members at the same feeder. To test whether these increased encounters led to changes in associations among the artificial groups, we examined social network metrics at a separate unrestricted feeder station using data from before and after the manipulation. Statistical analyses were performed in R (R Core Team, 2026) using the packages brms (Bürkner, 2017; Bürkner, 2018), emmeans (Lenth et al, 2026), igraph (Csárdi & Nepusz, 2006), assortnet (Farine, 2014), and asnipe (Farine, 2013).

To create artificial groups to manipulate, we first defined natural foraging associations before the manipulation using social network analysis from the unrestricted feeders’ data. For all the individuals in the colony networks, we calculated pairwise genetic relatedness using Wang’s estimate of relatedness (Wang et al., 2011) based on 16 microsatellites (Paquet et al., 2015). Dyads with relatedness estimates above 0.2 were classified as more related, likely representing close relatives (e.g. half-siblings, full-siblings, or parent-offspring pairs, aunts/uncles-nephews/nieces, or grandparents-grandchildren) while dyads with estimates below 0.2 were considered less related, including unrelated individuals or more distant relatives such as first cousins, and more distant cousins. Relatedness estimates were validated against a pedigree derived from a combination of video observations and genetic data (Silva et al., 2018).

We then constructed binary “inverted” genetic networks, in which more related dyads (r> 0.2) were not assigned an edge, and less related dyads (r < 0.2) were assigned an edge. This network structure allowed us to apply two clustering algorithms that grouped individuals by low relatedness edges. We first applied the Louvain community detection algorithm (Blondel et al., 2008) to optimize modularity to identify preliminary clusters of less genetically related individuals. Then, we applied hierarchical clustering to merge the Louvain clusters and reduce the total number of clusters to match the available feeders in each colony. In this step, clusters with the lowest average relatedness were iteratively merged to maintain the lowest level of genetic relatedness within the final clusters. Each resulting artificial cluster was assigned to a corresponding feeder at its colony’s allocated restrictive feeding station. This design enabled us to manipulate fine-scale association patterns while still allowing the full colony to forage and flock together at the same station with multiple feeders.

We tested whether the artificial clusters were less genetically related than the natural foraging groups. To do this, we used social networks from unrestricted feeders pre-experiment to define natural clusters of preferred foraging associations, applying the same Louvain and hierarchical clustering approach used for the artificial clusters. Both sets of clusters (foraging and genetic) had similar group sizes, and we compared the number of related edges in each. The natural foraging clusters contained significantly more related edges than the artificial clusters (Welch’s t-test: t = –3.44, p < 0.05, 95% CI [–0.20, –0.04]; mean related edge proportion: artificial = 0.043, natural = 0.165). Cluster sizes were similar between network types (mean: 17.5 individuals for artificial clusters vs. 17.3 for natural clusters). This indicates that the defined artificial clusters successfully reshuffled the kin-bias in the networks.

In the three sociable weaver colonies we manipulated, we assigned individuals to artificial clusters that matched the number of available selective feeders. Colonies 71 and 11 each had three feeders/clusters, and colony 8, being larger, had four. From the pre-manipulation networks, only two individuals did not have genetic data and were excluded, leaving 52 individuals in colony 71, 53 in colony 11, and 71 in colony 8. During the experiment, newly joined individuals (immigrants or fledglings from the 2024 breeding season) were added to the existing clusters (11 individuals were added to colony 71, 10 to colony 11, and 15 to colony 8). Individuals with genetic data were assigned to the cluster where they had the fewest related connections. Individuals that joined the colony without genetic data available could not be assigned to clusters based on relatedness. As newly captured birds, they were likely unrelated to existing colony members and were therefore assigned to clusters with the smallest group size (i.e. the least number of total individuals). This allowed us to balance cluster sizes with opportunities to form associations with less-related individuals. Final cluster sizes were: 63 individuals in colony 71 (24, 20, and 19 per cluster), 63 in colony 11 (21, 19, and 23), and 86 in colony 8 (26, 20, 21, and 19), with a total of 212 individuals.

#### (C) RFID-based feeding stations

Each sociable weaver colony was provided with their own feeding stations. Each colony had two feeding stations (unrestricted and restricted, Figure 1), located between 80 and 205m from the colony nests. Only one type of feeding station was open at a time, and when not used the other type of feeding station was concealed. Both stations consisted of three to four separate feeders, each with four RFID antenna perches and corresponding seed dispensers (Priority1 data loggers; Figure 1). Both feeding stations had similar perch distances (∼5cm) and provided the same canary seed mix.

Sociable weavers often forage together as a colony on insects and seeds (Maclean., 1973), but they can also feed in smaller groups or individually (Lloyd et al., 2017; Ferreira et al., 2020). The feeding stations were therefore designed to accommodate colony-level flocking behaviour when foraging, while allowing detection of fine-scale associations occurring at the individual feeders (Ferreira et al., 2020).

The first type of feeding station was made of unrestricted open feeders that allowed individuals to feed freely alongside others (Figure 1A & 1C; Ferreira et al., 2020). These feeders were used to collect data on foraging associations before the manipulation (January to April 2024), and after the manipulation (27^th^ November 2024 onward). Typically, the feeding stations were opened once every three days for a minimum of six hours. These data were used to detect any social association changes resulting from the experimental manipulation.

The second set of feeding stations consisted of restrictive experimental feeders, which we designed and built to manipulate foraging associations (Figure 1B). Each feeder was Raspberry Pi–based and programmed to recognize individual PIT-tag identities via antennas under the perches. Each feeder had four perches, each corresponding to a separate seed access point operated by an individual servo motor, allowing food access to be controlled at the perch level.

When a bird landed on a perch, the system read its unique RFID tag and then activated the relevant servo motors to control food access to all the perches depending on the stage of the manipulation and the identity of the tag (See stages below). To ensure that only the intended birds gained access to the food, the system continuously monitored the presence and identity of individuals on the perches, keeping food access open only while an authorized bird was detected in real time. The restrictive feeders were powered by large batteries (12V, 55AH) and solar panels, enabling them to run continuously throughout the experimental period.

The manipulation was conducted in three stages, designed to provide a prolonged habituation period and allow gradual adjustment to the feeding rules and restrictions. During all the manipulation phases, the unrestricted feeders were closed.

1. First, we ran a long **habituation phase** (April to August 2024) to allow individuals to adjust to using the new feeder designs, including the short dispensing delay and the sound of the servo motors. During this stage, if any PIT-tagged bird was detected at any feeder, food became available at all four perches, regardless of cluster membership.
2. Next, we ran an **association phase** (August to October 2024) to begin dividing the colony to feed with their associated artificial clusters. During this phase, each artificial cluster was assigned to one specific feeder. Only birds from the correct cluster could trigger food access at their assigned feeder. However, whilst any individual from that cluster was present, all perches at that feeder had food access, allowing any individual (regardless of cluster identity) to forage alongside them. At least one bird from the assigned cluster had to remain present for food access to continue. This stage encouraged birds to associate food access with their assigned feeder, and increased the likelihood of encountering other cluster members, whilst still permitting occasional foraging with prior social partners.
3. The final stage of the manipulation was the **restriction phase** (November 2024). During this phase, feeding became fully restricted to within clusters. Individuals could only dispense for themselves at their associated feeder, where food was provided only to the perch where the correct cluster individual was on, and no other birds could gain access unless they also belonged to that cluster. Therefore, individuals could feed alone or alongside any other members from their artificial cluster. This final stage reinforced cluster-based foraging encounters by limiting opportunities to feed with non-cluster individuals.

#### (D) Foraging social network construction

We constructed three foraging social networks for each colony: (1) the pre-manipulation networks from unrestricted feeders, (2) the during-manipulation networks from the restricted feeders, and (3) the post-manipulation networks from the unrestricted feeders (Figure 1).

The pre-manipulation networks (∼36 feeder data days collected between January and April 2024) provided baseline social associations before the manipulation. The during-manipulation networks were based on approximately 20 feeder days collected while individuals could only feed alone or with members of the same artificial cluster. Finally, the post-manipulation networks were constructed with unrestricted feeder data between November 2024 and February 2025 (∼32 feeder days). Immediately following the manipulation RFID data was collected intensively for seven to eight consecutive days, and then returning to regular collection patterns of once every three days. This sampling design provided a sufficiently large number of observations to construct robust social networks. To ensure reliable representation of individuals exposed to the manipulation conditions, only birds recorded on more than three days at the restrictive feeder during the manipulation were included in the post-manipulation network. To further reduce noise, individuals needed to be present for at least three separate feeder data days to be included in each social network.

Association strength in each of the networks were defined using a time-overlap method (Ferreira et al., 2020), where the time two individuals spent together at a single feeder was calculated as a simple ratio index (SRI). The SRI calculates the proportion of time that both individuals spent together relative to the total time each individual was observed at the feeding station (Cairns & Schwager, 1987). For the pre- and post-manipulation networks, any associations less than five seconds were excluded based on observations that most fights/displacements occur within this period (Ferreira et al., 2020). Removing these short overlaps increased confidence that associations longer than five seconds reflect tolerance rather than competition. This criterion also reduced noise in the networks and makes the resulting associations more meaningful. For the manipulation networks, this threshold was not applied because the RFID sensors at the restrictive feeders used a different reading pattern to continuously scan each perch.

#### (E) Statistical Analyses

During the manipulation, 95 individuals that were assigned to clusters used the restrictive feeders while access was limited to feeding alone or with members of the same artificial cluster (27 individuals from colony 11, 33 from colony 71, and 35 from colony 8).

To test whether the manipulation increased within-cluster foraging, we calculated weighted assortativity coefficients based on artificial cluster membership using the R package assortnet (Farine, 2014). Assortativity ranges from −1 (disassortative mixing) to 1 (complete assortative mixing), with higher values indicating stronger foraging associations between individuals belonging to the same artificial cluster. We expected assortativity to increase during the manipulation if birds preferentially associated with members of their assigned artificial cluster.

To quantify changes in pairwise foraging associations before and after the manipulation, we fitted a Bayesian zero-one inflated beta mixed model using the R package brms (Bürkner, 2017), which interfaces with Stan (Carpenter et al., 2017). Pairwise association strength (weighted time-overlap association index) was modelled as a function of experimental period (pre- or post-manipulation), dyad type (within or between artificial clusters), and their interaction. Random intercepts were included for dyad identity, colony, and individual identity using a multi-membership term for both members of each dyad. The model was fitted using four Markov chains with 5,000 iterations each (2,500 warm-up iterations) and the No-U-Turn Sampler (NUTS). Weakly informative normal priors were used for fixed effects and exponential priors for random-effect standard deviations.

Analyses were restricted to dyads present in both the pre- and post-manipulation social networks (n = 1,468). Dyads with no foraging association in either period (n = 182) were excluded because they had no opportunity to change their association strength, leaving 1,286 candidate dyads for analysis (875 between-cluster and 411 within-cluster dyads). Among the 411 within-cluster dyads, only fifteen pairs were closely related (relatedness > 0.2), providing limited variation to estimate the effect of relatedness reliably. Relatedness was therefore not included as a predictor.

Within-cluster dyads consisted of pairs in which both individuals belonged to the same artificial cluster and were permitted to feed together during the manipulation. Between-cluster dyads consisted of pairs from different artificial clusters and therefore could not feed together under the restrictive treatment. We quantified changes only for within-cluster dyads that experienced frequent encounters at the restrictive feeders and therefore were sufficiently exposed to the manipulation conditions.

To quantify exposure to the manipulation, RFID data collected at the restrictive feeders were used to identify flocks using the Psorakis co-occurrence algorithm (Psorakis et al., 2012), applied separately for each colony, feeder, and day. We then counted the number of flocks in which both individuals of each dyad were observed together, representing their opportunity to interact under the experimental conditions. Because the restrictive phase of the experiment was relatively short, we retained only within-cluster dyads that co-occurred in at least 95 manipulation flocks (n = 23), thereby ensuring substantial exposure to the manipulation. Control dyads consisted of between-cluster pairs for which both individuals were observed at least once at the restrictive feeders (n = 468). The final model therefore included 491 dyads each represented in both periods (982 observations).

Model convergence was good (all ^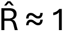^), with no divergent transitions and adequate effective sample sizes. Posterior predictive checks indicated good agreement between observed and simulated data, including both zero values and non-zero association strengths.

